# Beyond comparison: Brillouin microscopy and AFM-based indentation reveal divergent insights into the mechanical profile of the murine retina

**DOI:** 10.1101/2024.01.24.577013

**Authors:** Stephanie Möllmert, Marcus Gutmann, Paul Müller, Kyoohyun Kim, Jana Bachir Salvador, Serhii Aif, Lorenz Meinel, Jochen Guck

## Abstract

Mechanical tissue properties increasingly serve as pivotal phenotypic characteristics that are subject to change during development or pathological progression. The quantification of such material properties often relies on physical contact between a load-applying probe and an exposed sample surface. For most tissues, these requirements necessitate animal sacrifice, tissue dissection and sectioning. These invasive procedures bear the risk of yielding mechanical properties that do not portray the physiological mechanical state of a tissue within a functioning organism. Brillouin microscopy has emerged as a non-invasive, optical technique that allows to assess mechanical cell and tissue properties with high spatio-temporal resolution. In optically transparent specimens, this technique does not require animal sacrifice, tissue dissection or sectioning. However, the extent to which results obtained from Brillouin microscopy allow to infer conclusions about potential results obtained with a contact-based technique, and *vice versa*, is unclear. Potential sources for discrepancies include the varying characteristic temporal and spatial scales, the directionality of measurement, environmental factors, and mechanical moduli probed. In this work, we addressed those aspects by quantifying the mechanical properties of acutely dissected murine retinal tissues using Brillouin microscopy and atomic force microscopy (AFM)-based indentation measurements. Our results show a distinct mechanical profile of the retinal layers with respect to the Brillouin frequency shift, the Brillouin linewidth and the apparent Young’s modulus. Contrary to previous reports, our findings do not support a simple correlative relationship between Brillouin frequency shift and apparent Young’s modulus. Additionally, the divergent sensitivity of Brillouin microscopy and AFM-indentation measurements to cross-linking or changes *post mortem* underscores the dangers of assuming both methods can be generally used interchangeably. In conclusion, our study advocates for viewing Brillouin microscopy and AFM-based indentation measurements as complementary tools, discouraging direct comparisons *a priori* and suggesting their combined use for a more comprehensive understanding of tissue mechanical properties.

## 1. Introduction

Retinal pathologies, such as age-related macular degeneration (AMD) and hereditary retinal dystrophies, are responsible for a considerable portion of untreatable blindness worldwide^[1]^. Despite substantial efforts directed toward elucidating their etiology, there exists a noteworthy gap in investigating the mechanical properties of ocular tissues affected during typical human development and aging. Mechanical tissue properties have emerged as phenotypic markers contributing to a comprehensive understanding of cellular and tissue physiology in both developmental and pathological contexts. A more profound comprehension of the mechanical characteristics of retinal tissues holds the potential to yield novel insights into pathogenic mechanisms and may offer direction towards innovative or enhanced therapeutic approaches.

The measurement of mechanical tissue properties is predominantly assessed by exerting an external force, recording the resulting deformation, and subsequently extracting a proportionality factor that equates to an intrinsic material property which describes the material’s resistance to the deformation. Atomic force microscopy (AFM)-based indentation measurements, for instance, utilize a flexible cantilever with a geometrically well-defined probe on its tip to indent the sample with a preset force^[2,3]^. As the cantilever approaches the sample, its deflection is recorded as a function of distance from the sample surface. Upon reaching the surface, it pushes into the sample until the preset force is reached. This yields a force-distance curve in which the slope of the indentation segment is indicative of the material’s resistance to deformation. Due to this operation principle, indentation measurements necessitate almost always the preparation of *ex vivo* samples to enable the physical contact between the probe and the sample surface. The resistance to deformation is termed Young’s modulus, and has been shown to change in response to central nervous system (CNS) injury^[4,5]^, during CNS development^[6]^ and aging^[7]^, and upon changes of extracellular matrix composition in the CNS^[8,9]^. As other tissues of the CNS^[4,7,10]^, the retina has been described as heterogeneous with respect to local tissue architecture and composition^[11]^, and concurrent mechancial properties^[12,13]^. Ocular structures, such as ocular surface, lens, cornea, mucins, Bruch’s membrane as well as isolated retinal cells have been studied using AFM^[14]^. In fact, AFM-based indentation measurements have been employed to investigate the stiffness of the retinal microvasculature^[15]^ and the inner retinal surface^[13]^. Yet, little attention has been directed toward a systematic analysis of the mechanical properties of the individual retinal layers.

Non-invasive methods, such as ultrasound elastography (USE), magnetic resonance elastography (MRE) or optical coherence elastography (OCE), have been used to investigate ocular tissues^[1]^ *in vivo*, but face limits due to insufficient resolution (MRI)^[16]^, or inaccessibility of the targeted tissue (USE, OCE) ^[1,17,18]^, as well as concomitantly damagingly high intensities for mechanical excitation (OCE)^[17]^.

Contrastingly, *ex vivo* measurements on dissected retinae using both OCE and Brillouin microscopy have been able to successfully characterize individual retinal layers from fresh and chemically fixed retinal cups^[19]^. Brillouin microscopy is a quantitative imaging modality that detects an inelastic scattering process of light from thermally induced density fluctuations^[20]^. The density fluctuations are dependent on the material properties of the measurement volume, *i*.*e*. the longitudinal modulus, the viscosity, and the density. The longitudinal modulus is linked to the change of frequency between incident and scattered light (Brillouin frequency shift). The viscosity of the material contributes to the attenuation of the acoustic wave propagation due to energy dissipation, and scales with the linewidth of the Brillouin scattered light in a spectrum^[20]^. Brillouin microscopy therefore allows to optically, non-invasively probe viscoelastic properties of complex, transparent biological samples *in vivo*, but to some extent also more opaque materials with limited penetration depth *ex vivo*.

Despite the fact that AFM-indentation and Brillouin microscopy quantify material parameters that occur on vastly different time- and length-scales, Brillouin microscopy has often been compared to AFM-based measurements. Although phenomenological correlations between longitudinal modulus (or Brillouin frequency shift) and apparent Young’s modulus have been reported^[21-23]^, it is currently unclear to what extent we can infer information about one modulus from measuring the respective other. Therefore, it has been suggested that these correlations should be calibrated beforehand and for each sample^[24]^.

In this work, we systematically quantify the mechanical properties of acutely sectioned murine retina tissues using both AFM-based indentation and Brillouin microscopy. Our results reveal a distinctive mechanical profile in the retinal layers concerning the Brillouin frequency shift, Brillouin linewidth, and apparent Young’s modulus, which deviates from previous reports on specific retinal layers^[23]^. Contrary to previous reports^[21-23]^, our findings do not support a straightforward correlative relationship between the Brillouin frequency shift and the apparent Young’s modulus. To augment our investigations, we evaluate layer-specific mechanical responses to genipin-mediated cross-linking—a non-cytotoxic compound derived from the *Gardenia jasminoides* plant known for its ability to induce chemical cross-linking of polymers with amino groups, such as collagen^[25-29]^. During the treatment of ocular pathologies, genipin has been reported to effectively induce scleral stiffening *in vivo* and *ex vivo* rendering it a biocompatible candidate for clinical translation^[29]^. Additionally, we explore mechanical changes with an increased *post mortem* time interval. The disparate sensitivity of Brillouin microscopy and AFM-indentation measurements to these alterations emphasizes the dissimilar nature of each method. In conclusion, our results advocate for the combined use of both methods for a more comprehensive mechanical characterization, discouraging *a priori* assumptions of mutual verification.

## 2. Method

### 2.1 Animal husbandry and handling

Husbandry and experimental procedures were performed in accordance with the Association for Research in Vision and Ophthalmology Statement for the Use of Animals in Ophthalmic and Vision Research. This study exclusively used C57BL/6JRj mice that were bred and kept in the animal facility of the University of Würzburg and the Biologisch-Technisches Entwicklungslabor Erlangen. All animals were between three months and six months of age, and euthanized by cervical dislocation. All measurements started between 20 minutes and 90 minutes *post mortem*, depending on treatment.

### 2.2 Retinal tissue dissection

Eyes were removed immediately after cervical dislocation, with 45° angled forceps and, to prevent desiccation, transferred to a tissue culture medium (TCM) that contained (in mM) 136 NaCl, 3 KCl, 1 MgCl_2_, 10 HEPES, 2 CaCl_2_ and 11 glucose, adjusted to pH 7.4 with NaOH. The dissection of the immersed retina was performed in the lid of a 35 mm petri dish. First, muscle and connective tissue were removed as thoroughly and gently as possible using forceps (Dumont #5, Fine Science Tools GmbH, Heidelberg, Germany) and scissors (2.5 mm Cutting Edge Vannas Spring Scissors, Fine Science Tools GmbH, Heidelberg, Germany). Then, the eye was held at the limbus using forceps to facilitate a minimal incision with a cannula. Thereafter, the cornea was removed with spring scissors after circular cuts along the *Ora Serrata*. The exposed lens was separated and the vitreous was detached with forceps (Dumont #5-45, Fine Science Tools GmbH, Heidelberg, Germany). The optic nerve was cut at the location between the retina and the eyecup to ease the separation of the retina. Finally, a small incision was made between the retina and the sclera-choroidea in the direction of the optic nerve, so that the sclera-choroidea could be opened with the help of two forceps to detach the retinal cup as a whole.

### 2.3 Tissue embedding and sectioning

Dissected retinal cups were immersed in liquid low-gelling-point agarose (2.5% in TCM, cooled to 31°C, Sigma-Aldrich, A0701) in a 3 mm diameter cell culture dish. The cups were centered and positioned with insect pins, so that the cup openings were facing the dish wall. Upon solidification, a piece of agarose gel containing the retinal cup was cut out into a block of approximately 1 cm x 1 cm x 1 cm, and was sectioned with an oscillating-blade vibratome while the cup opening was facing the experimenter. Most precise and even tissue sectioning was achieved by using a cutting frequency of 100 Hz, a velocity of 2.5 mm/s and an amplitude of 0.4 mm. A section thickness of 250 μm has proven to be optimal for subsequent sample mounting. Since our experiments comprised the investigation of temperature changes during the preparation procedure, we used a buffer temperature of approximately 15 – 18 °C for measurements following a flat temperature profile during tissue preparation and 4 – 6 °C for measurements following a preparation procedure that involved incubation in ice-cooled medium. This procedure yielded acute retinal sections in which the individual retinal layers were exposed and could then be accessed either with the indenter tip or optically with the laser. All measured tissue sections originated from a plane close to the optic nerve and displayed minimal curvature. The acute retinal tissue sections were incubated in TCM at room temperature or on ice, as indicated, until further processing for mechanical measurements. All experiments that involved the addition of genipin (10 mM in 1x TCM, Sigma Aldrich, G4796) required a change of the glucose content of the TCM to 5 mM and an incubation time of 90 minutes prior to measurements. The control experiments for the genipin treatments were performed with TCM with 5 mM glucose and the same incubation time, but without genipin.

### 2.3 Tissue section mounting

Retinal sections were immobilized on culture dishes with Histoacryl® (B. Braun, 9381104) which was sparsely applied between the bottom of the dish and the agarose embedding, but far from the tissue section. For indentation measurements, the sections were immobilized on tissue culture plastic (TCP). For Brillouin microscopy, the sections were immobilized on glass bottom dishes. To prevent the section from floating upwards and changing the penetration depth and defocusing during Brillouin microscopy, glass cover slips were installed above the tissue section and held in place by putty at the plastic rim of the glass-bottom dish thereby serving as a roof against which the tissue section might float without exerting pressure on the tissue itself. The tissue sections were submerged in TCM during all measurements.

### 2.4 Brillouin microscopy

Brillouin microscopy was performed using a custom-built confocal Brillouin microscope based on a two-stage virtually imaged phase array (VIPA). The detailed working principle is described in ^[21]^. Briefly, we used a frequency-modulated diode laser beam (*λ* = 780.24 nm, DLC TA PRO 780, Toptica Photonics AG, Germany) as an illumination source. The central frequency of the beam was stabilized to the D_2_ transition of ^85^Rb by monitoring the intensity of the probe beam after it passed through a rubidium molecular absorption cell (TG-ABRB-I85-Q, Precision Glassblowing Inc., USA). In order to suppress the amplified spontaneous emission (ASE) and clean the laser spectrum, we implemented a Fabry-Pérot interferometer (FPI) in the illumination path with the cavity length of a piezo-tunable etalon (OP-1986-102, LightMachinery Inc., Canada) stabilized to the central laser frequency. The FPI led to a reduction of the laser mode to ASE ratio by 20 to ∼85 dB, and the ASE was further suppressed by a monochromatic detector (DFC MD, Toptica). The laser beam was then coupled into a single-mode fiber and guided to a custom-built inverse confocal microscope. The confocal microscope was based on an inverted microscope stand (Axio Observer 7, Carl Zeiss AG, Germany). The laser beam was focused in the tissue sample by an objective lens (NA 0.5, 20×, EC Plan-Neofluar, Carl Zeiss) yielding a resolution of 1 μm in lateral and 5 μm in axial direction. Sample positioning and translation during the scanning process was facilitated by a motorized stage using a step size of 1 μm in *x*- and *y*-direction for all maps. The penetration depth of the laser beam inside the tissue sample was the maximum distance from the cover slip at which the live spectra was still clearly obtainable. Depending on treatment, typical penetration depth ranged from 30 μm to 60 μm. In order to maintain the viability of the specimen, the intensity of the focused laser beam on the specimen was kept at 10 mW. The backscattered light was collected by the same objective lens and coupled into a second single-mode fiber, which achieved confocal sectioning. The fiber-coupled light was guided to the Brillouin spectrometer. In the spectrometer, the light was collimated and passed through the same rubidium molecular absorption cell for frequency stabilization for attenuating the Rayleigh (elastic) scattered and reflected light. Then, the beam was guided to two-stage VIPA etalons (OP-6721-6743-4, LightMachinery) with the 15.2 GHz free spectral range. The VIPA etalons led to angular dispersion of light with different frequencies^[21]^, which spectra were recorded by a scientific complementary metal-oxide-semiconductor camera (Prime BSI, Teledyne Photometrics, USA) with an acquisition time of 0.25 sec per spectrum and a 2×2 binning. For each pixel, two spectra were recorded and averaged. The acquisition of Brillouin microscopy images and the control of the microscope were facilitated by a custom-made software written in C++. The source code can be found at https://github.com/BrillouinMicroscopy/BrillouinAcquisition. From the recorded images, the one-dimensional Brillouin spectra were extracted, and the position, width, and intensity of the Brillouin and Rayleigh peaks were determined by fitting Lorentzian functions to every peak. To calibrate the frequency axis, a reference measurement of methanol- and water-filled cuvettes placed in the calibration beam path was acquired every 10 min by moving a motorized mirror into the calibration beam path. This allowed us to obtain an absolute value for the Brillouin frequency shift and compensated for possible drifts of the VIPA setup during the measurement. Acquired data were analysed using the custom-made Python software BMicro 0.8.1. The source code is available at https://github.com/BrillouinMicroscopy/BMicro.

To be able to allocate Brillouin frequency shift and Brillouin linewidth values to distinct retinal layers in an unbiased, orthogonal manner, we used Impose 0.4.1, a custom-made software written in Python that can be found at https://github.com/GuckLab/impose). To this end, all tissue sections were chemically fixed with 4% PFA in PBS overnight upon Brillouin microscopy data acquisition, and subsequently washed and stained with DAPI. This was followed by confocal fluorescence microscopy using a Plan-Apochromat 20×/0.8 objective on a Zeiss LSM 980 confocal microscope. Impose then allowed to superimpose both imaging modalities, Brillouin microscopy and fluorescence microscopy images, to match Brillouin frequency shifts and linewidths with correct retinal layers based on their DAPI signal or lack thereof. We acquired two spectra for each pixel, whereas the total amount of pixels for each layer corresponded to approximately 400 to 2000 different locations, depending on the layer thickness. The Brillouin frequency shift and linewidth values extracted for one layer from one retina were averaged and contributed one datum to the final plot.

Some Brillouin microscopy measurements were acquired on two identical, but physically and spatially separate setups. This allowed us to minimize the total amount of animals while still increasing the sample size. Despite their identical configuration and calibration samples, the two setups yielded different Brillouin frequency shifts and linewidths for the same water sample. To account for this discrepancy, we acquired Brillouin microscopy measurements of the same water sample on days when we employed both setups (Supplementary Figure 1). The respective daily offsets in Brillouin frequency shifts and linewidths were then corrected by adding the difference to the values acquired with the second setup. This procedure was necessary for the control measurements for the genipin treatment.

To calculate the longitudinal modulus from the Brillouin frequency shift, the following equation applies:

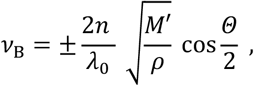

where *ν*_B_ denotes the Brillouin frequency shift, *n* is the refractive index, *λ*_0_ is the wavelength of the incident light, *M’* is the longitudinal storage modulus, *Θ* is the scattering angle, and *ρ* is the mass density of the material in the measured volume. The viscosity *η* of the material is linked to the Brillouin linewidth *Δν*_B_ by:

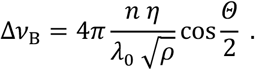

In Brillouin microscopy, mechanical and optical material properties are coupled^[20]^. In order to calculate the longitudinal modulus and the viscosity present in the measurement volume, the refractive index and the density of that volume must be known. Since direct measurements of refractive index and density, especially for extended tissues, are not straightforward^[30]^, Brillouin frequency shift and Brillouin linewidth are reported as proxies for longitudinal modulus and the viscosity in this study, respectively.

### 2.4 AFM-based indentation measurements

Indentation measurements enabled by atomic force microscopy (AFM) and simultaneous brightfield microscopy was performed with the NanoWizard 4 (JPK Instruments/Bruker, Berlin) and the upright Axio Zoom.V16 stereo microscope with a PlanApo Z 0.5x objective (Carl Zeiss Microscopy, Jena). For indentation experiments, polystyrene beads (d = (10.03 ± 0.14) μm, Microparticles GmbH, PS-R-10.0) were glued to tipless silicon cantilevers (Arrow-TL1, NanoWorld) with epoxyglue. Cantilevers were calibrated using the thermal noise method ^[31]^ prior to experiments. Only cantilevers with spring constants between 0.030 N/m and 0.045 N/m were used.

To obtain detailed spatial information about the mechanical properties of the individual layers of the murine retina, all indentation measurements were carried out on acute sections that spanned the entire diameter of the retinal cup and displayed minimal curvature to exclude contribution of underlying structures from neighboring layers. Simultaneous brightfield microscopy with high intensity transmitted light was sufficient to identify individual layers and enabled visual guidance during the manual positioning of the indenter tip. Each layer was indented once in approximately 30 to 100 different locations. The indentation force was 4 nN and the indentation speed was 10 μm/s. All indentation measurements took place at 21°C room temperature. All force-distance curves were analyzed using a custom-designed Python software PyJibe 0.14.3 (https://github.com/AFM-analysis/PyJibe). Here, the approach segment of each indentation curve is fitted with the Hertz model modified for a spherical indenter:

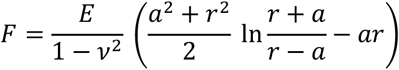

with

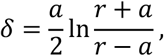

where *F* denotes the indentation force, *δ* the indentation depth, *r* the indenter radius, and a is the radius of the circular contact area between indenter and sample ^[32,33],34^. The Poisson’s ratio *ν* was set to 0.5 for all analyses. The Young’s modulus *E* was used as the fitting parameter and served as a measure for the apparent elastic resistance of the probed sample to deformation. Since both measurement and analysis described in this part of the study approximate the tissue as a purely elastic solid and do not account for viscous material properties, the values of the Young’s modulus are termed ‘apparent’.

### 2.5 Data visualization and statistical analysis

The graphical data representation was compiled with Python and utilizes in most cases raincloud plots^[35]^ in which the individual data points are means from the measurement data for each sample. The boxplot shows the median with the interquartile range (IQR), and whiskers that extent to the most extreme data points that lie within 1.5 IQR. The half-violin plot displays the kernel density estimate calculated using a bandwidth *h* with

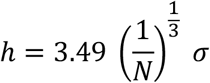

where *N* denotes the sample size and *σ* denotes the sample standard deviation ^[36]^. Statistical analyses were performed in Origin (OriginLab, OriginPro 2021b). *N* refers to the number of sections. Whenever multiple sections were obtained from the same retina, these sections were distributed over different experiments and measurement methods. All data were subjected to normality tests using the Chen–Shapiro test. Since almost all normality test methods perform poorly for small sample sizes (less than or equal to 30), and Chen– Shapiro test results additionally yielded too few data points to assume normality, we proceed with non-parametric statistical tests for all analyses. In particular, the data in Figure 1 comprises independent samples. Here, we employed the Kruskal–Wallis ANOVA which yielded for subplot b) a test statistic *χ*^**2**^(5)=55.915 and a probability *p*=8.461*10^-11^ with *N*=21, for subplot c) *χ*^**2**^(5)=41.751 and *p*=6.616*10^-8^ and *N*=21, and for subplot d) *χ*^**2**^(5)=19.673 and *p*=0.001 with *N*=6. Dunn’s *post hoc* analysis revealed a statistically significant difference between the indicated retinal layers. Exact p-values are denoted in the respective positions in the graphs. The Kruskal-Wallis effect size *η*^2^ was calculated using *η*^2^ with

**Figure 1:**
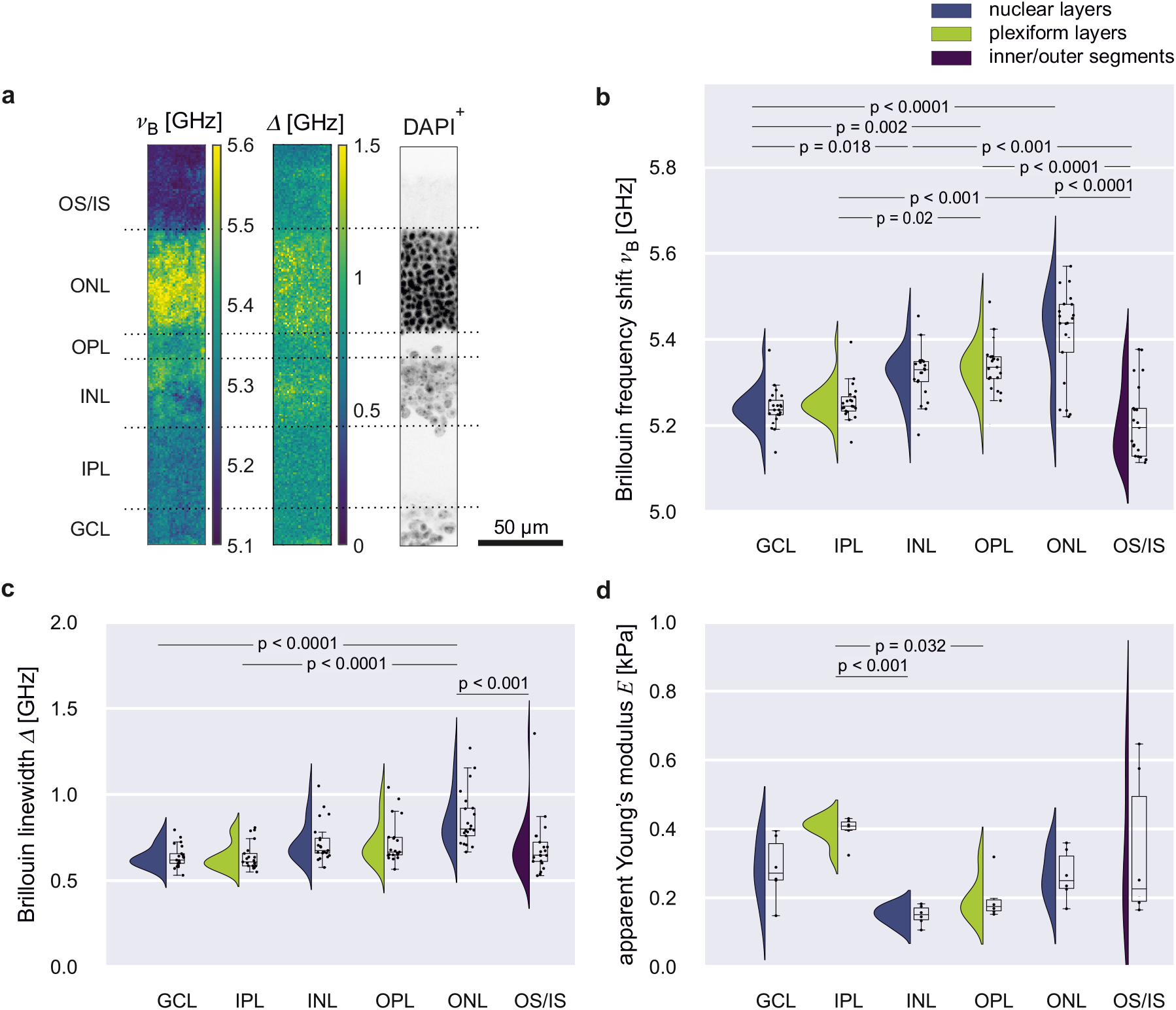
The mechanical phenotype of acutely sectioned retinal tissues. **a)** Brillouin microscopy maps display a layered distribution of both the Brillouin frequency shift ν_B_ and Brillouin peak linewidth Δ. All section were subjected to a DAPI stain after the measurement. DAPI-positive (DAPI+) regions indicate the nuclear layers in the retina, i.e. the ganglion cell layer (GCL), the inner nuclear layer (INL), and the outer nuclear layer (ONL). DAPI-negative regions correspond to the inner (IPL) and the outer plexiform layer (OPL), as well as the outer and inner segments (IS/OS). The superposition of both the Brillouin microscopy maps and the fluorescence image facilitates the unbiased allocation of the Brillouin microscopy values to distinct retinal layers. For each pixel two spectra were recorded, whereas the total amount of pixels for each layer corresponded to approximately 400 to 2000 different locations in the respective retinal layer. The Brillouin frequency shift and linewidth values extracted for one layer from one retina were averaged and contributed one datum to the final plot. This procedure was conducted for 21 retinae. **b)** The distribution of the Brillouin frequency shift means across individual retinal layers (N=21 retinae). **c)** The distribution of the Brillouin linewidth means across individual retinal layers (N=21 retinae). **d)** The distribution of the apparent Young’s modulus means of individual retinal layers obtained with indentation measurements. Each layer was indented once in approximately 30 to 100 different locations. Indentation measurements for one retinal layer and retina were then averaged and contributed one datum in the final plot. This procedure was conducted for six retinae (N=6 retinae). Graphs display raincloud plots with half violins showing the data distribution. Box plots showing the median, interquartile range (IQR) and whiskers for minimum and maximum data points. The scattered data points show all means including outliers (>1.5 IQR). P-values indicate significance level of difference between retinal layers.

**Figure 2:**
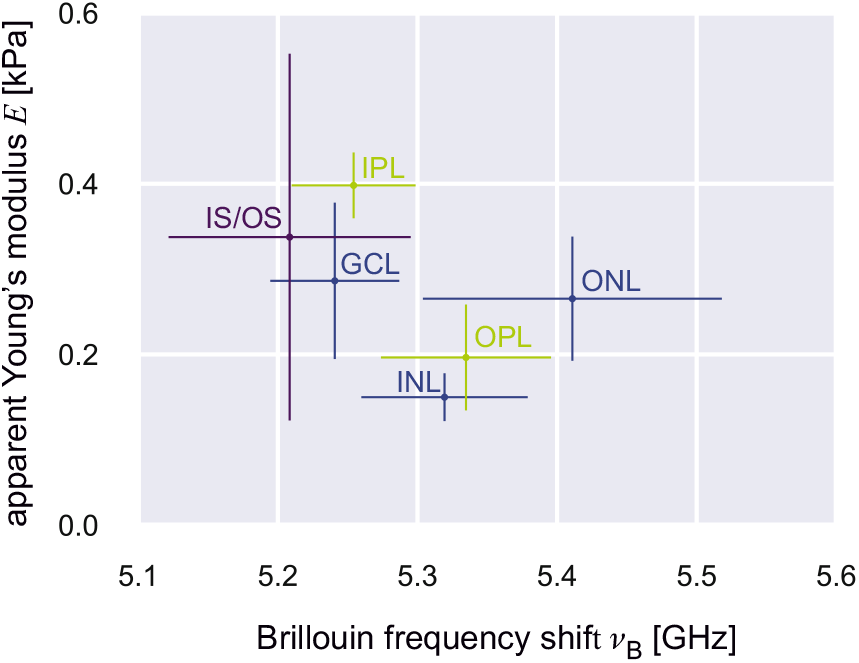
Correlative analysis of the Brillouin frequency shift and the apparent Young’s modulus. The mean values for the Brillouin frequency shift (*N*=21) and the apparent Young’s modulus (*N*=6) obtained from unaltered acute retina sections do not display a simple correlative relationship. Dots indicate mean values and whiskers show the standard deviation. Data taken from Fig.1 b) and d).

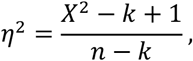

where *χ*^**2**^ is the Kruskal–Wallis test statistic value; *k* is the number of groups and *n* is the total number of observations^[37]^. In our case *η*^2^=0.42 (large effect)^[38,39]^ for b), *η*^2^=0.31 (large effect) for c), and *η*^2^=0.49 (large effect) for d). The data in Figure 3 comprises independent samples that are to be compared pairwise, each treated layer to its control measurement. Therefore, we used the Mann–Whitney test which yields a test statistic *U*, a *Z* score and a probability *p* for each pairwise comparison. The effect size *r* was calculated using

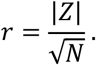

**Figure 3:**
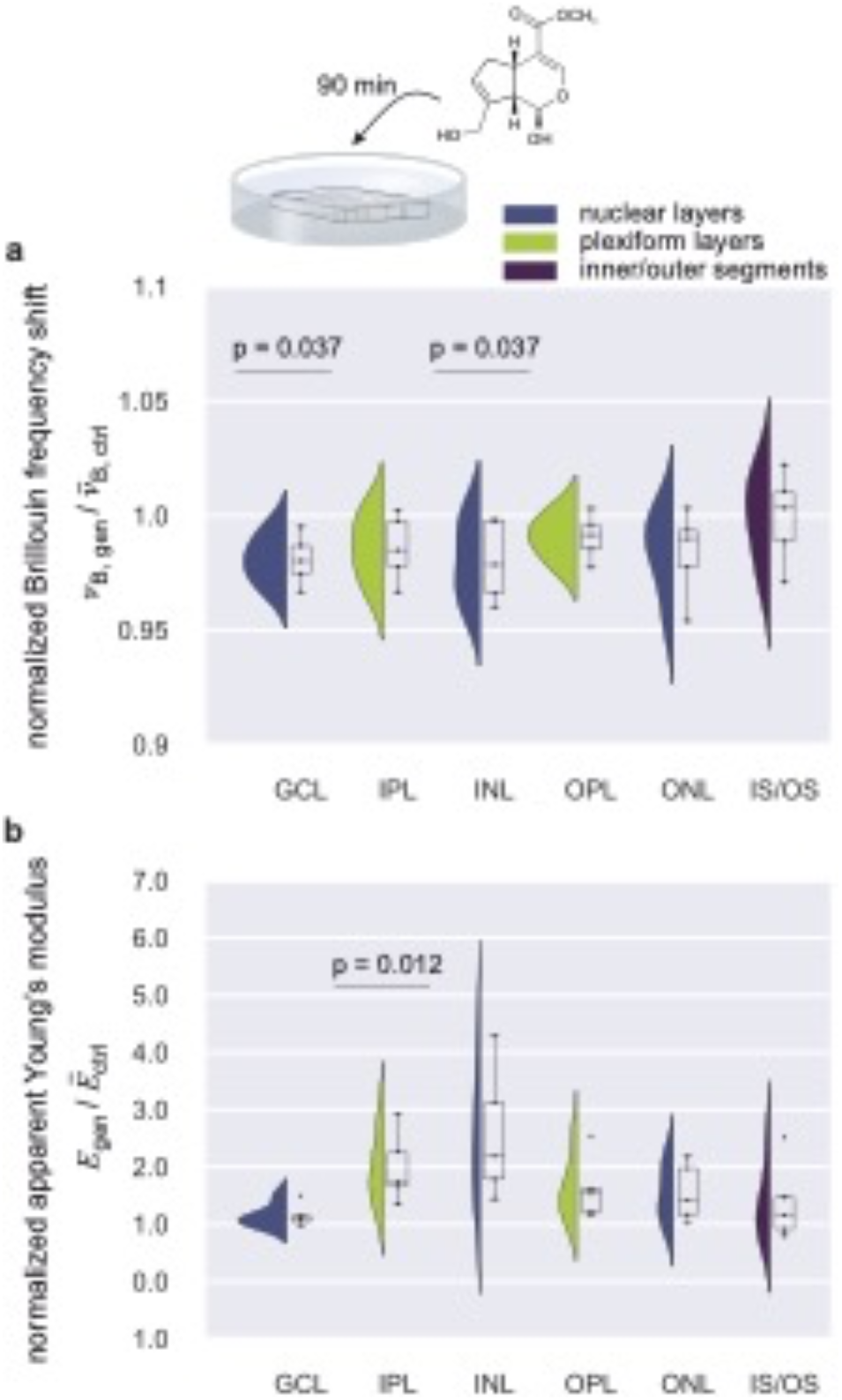
The impact of genipin cross-linking on the mechanical properties of the retina. **a)** The Brillouin frequency shift (*N*=5) and **b)** the apparent Young’s modulus of genipin treated retina sections (*N*=5) normalized to layer means from respective control measurements (*N*=5 each). Graphs display raincloud plots with half violins showing the data distribution. Box plots showing the median, interquartile range (IQR) and whiskers for minimum and maximum data points. The scattered data points show all means including outliers (>1.5 IQR). P-values indicate significance level of difference between retinal layers.

The analysis for both the GCL and the INL in Figure 3 a) yielded *U*=23, *Z*=2.089 and *N*=5 which results in *r*=0.93 (large effect) for each comparison. In subplot c), the comparison for the IPL yielded *U*=0, *Z*=-2.507 and *N*=5 which results in *r*=1.12 (large effect). The analysis for the timelapse experiments in Figure 4, yielded for the Brillouin frequency shifts in b) for the OPL *χ*^**2**^(6)=10.147, *p*=0.119, *N*=3, and *η*^2^=0.296 (large effect); for the ONL *χ*^**2**^(6)=4.207, *p*=0.649, *N*=3, and *η*^2^=0.128 (intermediate effect); and for the IS/OS *χ*^**2**^(6)=1.385, *p*=0.967, *N*=3, and *η*^2^=0.33 (large effect). Regarding the linewidths in Figure 4c), we calculated for the INL *χ*^**2**^(6)=2.511, *p*=0.867, *N*=3, and *η*^2^ = 0.249 (large effect); for the OPL *χ*^**2**^(6)=2.165, *p*=0.904, *N*=3, and *η*^2^ = 0.274 (large effect); for the ONL *χ*^**2**^(6)=1.368, *p*=0.968, *N*=3, *η*^2^=0.331 (large effect); and for the IS/OS *χ*^**2**^(6)=2.32, *p*=0.888, *N*=3, and *η*^2^ = 0.263 (large effect). Figure 4 d) comprises multiple independent samples, therefore a Kruskal–Wallis ANOVA yielded *χ*^**2**^(2)=6.489 and a probability *p*=0.034 with *N*=3. Dunn’s *post hoc* analysis revealed a statistically significant difference only between 0.5 hours *post mortem* (hpm) and 6 hpm in the INL. The effect size *η*^2^ = 0.75 (large effect).

**Figure 4:**
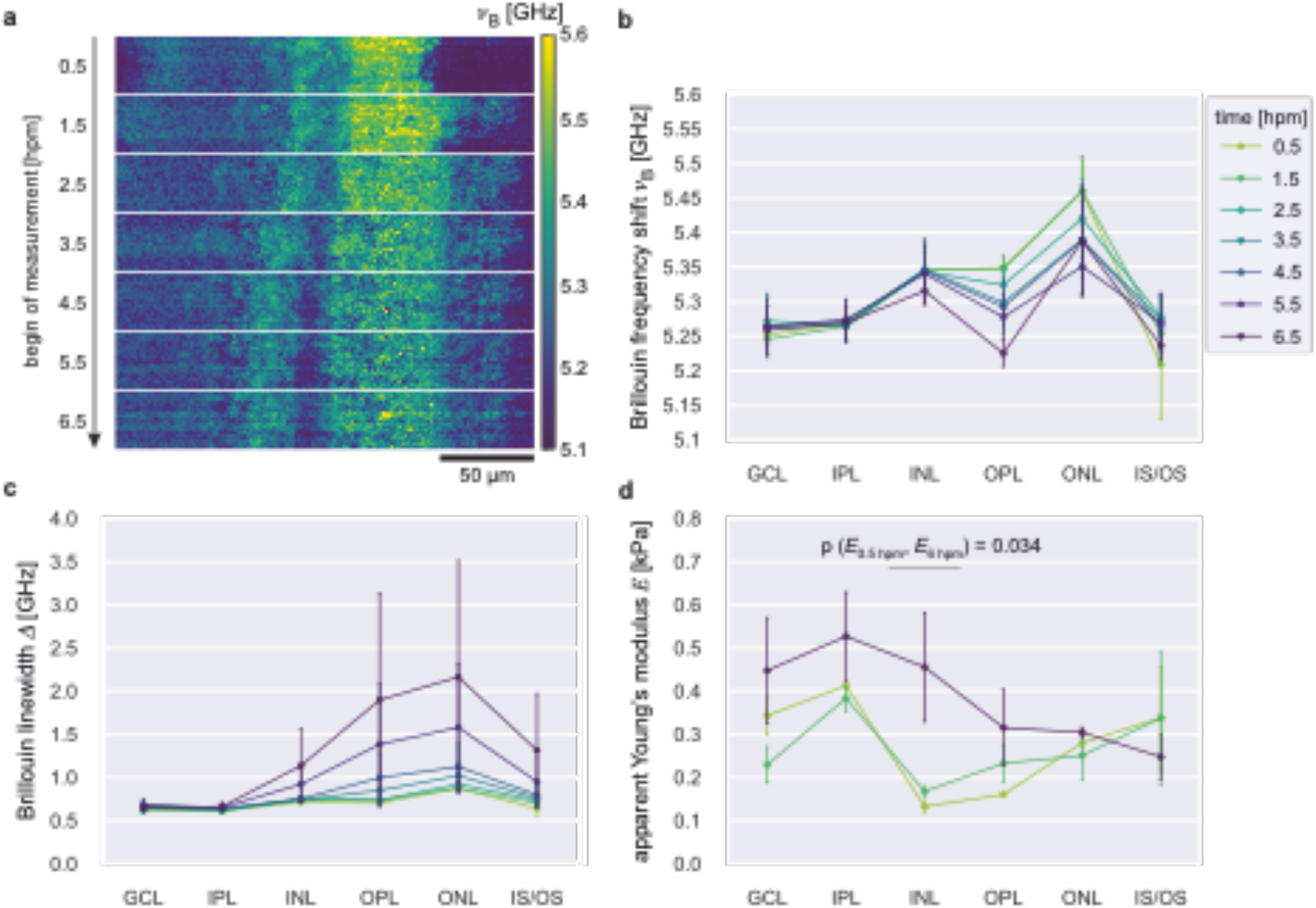
Mechanical properties change post mortem. **a)** Brillouin microscopy maps showing the temporal evolution of the Brillouin frequency shift of the same retina section. **b)** The Brillouin frequency shift profile for all retinal layers averaged for *N*=3 specimens over time. **c)** Brillouin linewidth profile for all retinal layers averaged for *N*=3 specimens over time. d) The mean apparent Young’s modulus of *N*=3 specimens for each layer at 0.5 hpm, 1.5 hpm and 6 hpm. Pointplots show the connected means for each layer with the standard error of the mean.

## 3. Results

### 3.1 Retinal layers display distinct mechanical properties

To systematically quantify the mechanical properties of the murine retina, we implemented a meticulous preparation protocol facilitating the concurrent application of confocal Brillouin microscopy and AFM-based indentation measurements on the same specimen. Confocal Brillouin microscopy yielded maps that displayed a stratified distribution of both Brillouin frequency shift values *ν*_B_ and linewidth values *Δ* that were reminiscent of the stratified structural organization of the retinal tissue (Figure 1a). To allocate the spectra obtained with Brillouin microscopy to distinct retinal layers in an unbiased manner, all sections were chemically fixed, incubated with the nuclear stain DAPI and imaged using a confocal fluorescence microscope after the Brillouin microscopy measurement. In fluorescence images, DAPI-positive (DAPI+) regions indicate the nuclear layers in the retina, *i.e*. the ganglion cell layer (GCL), the inner nuclear layer (INL), and the outer nuclear layer (ONL) (Figure 1a). DAPI-negative regions correspond to the inner (IPL) and the outer plexiform layer (OPL), as well as the outer and inner segments (IS/OS). The overlay of Brillouin microscopy maps onto the retinal layers identified by fluorescence microscopy aided by custom software (Impose, see 2.4 Brillouin microscopy) facilitates an unbiased allocation of the Brillouin microscopy values to distinct retinal layers and revealed unique Brillouin frequency shift and linewidth profiles for each layer. For each specimen, all spectra were averaged for each retinal layer and contributed with one mean value to the final sample size. Notably, from the innermost GCL to the ONL, both Brillouin frequency shift and linewidth exhibited increasing mean values, with the IS/OS region displaying significantly lower values (Figure 1b). This trend was more pronounced for the Brillouin frequency shift and subtler for the linewidth (Figure 1c). To complement the contact-free Brillouin microscopy, we employed AFM-based indentation measurements guided by simultaneous brightfield microscopy for precise layer targeting. The quantification of the apparent Young’s modulus of individual retinal layers shows that the GCL and the IPL exhibited the highest resistance to compression, while the INL, OPL and ONL displayed comparatively lower values (Figure 1d). The IS/OS layer demonstrated a resistance profile comparable to GCL and ONL, but with broader distribution (Figure 1d).

#### 3.1.1 Brillouin frequency shift and apparent Young’s modulus do not correlate

Using acutely sectioned retinal tissues ensured uniform and tightly controlled environmental conditions, and consistent measurement/detection directionality for both methods. This facilitated an exploration into the impact of the different time- and length-scales on which both methods operate, enabling a direct comparison of Brillouin frequency shifts and apparent Young’s moduli. Contrastingly to previous reports, a correlative analysis reveals that the data sets obtained with Brillouin microscopy and AFM-based indentation display no correlation (Figure 2).

### 3.2 Cross-linking with genipin induces divergent mechanical changes in distinct retinal layers

To investigate the impact of chemical cross-linking on both Brillouin microscopy and AFM-based indentation measurements, we subjected freshly sectioned retinal tissues to incubation with genipin. Concurrently, control measurements were conducted on tissue sections subjected to identical preparation and incubation conditions, excluding the application of genipin. The measurements obtained from genipin-treated sections were normalized with respect to the layer-specific means derived from those control experiments. Brillouin microscopy analyses revealed a significant reduction in the Brillouin frequency shift (Figure 3a) exclusively within the GCL and INL following genipin-mediated cross-linking, while linewidth values remained unaffected (Supplementary Figure 2). AFM-indentation measurements on these samples indicated a significantly elevated apparent Young’s modulus for the IPL and INL (Figure 3b). When genipin was applied to the retinal cup before sectioning, the Brillouin frequency shift of the ONL was significantly reduced (Supplementary Figure 3a), and the apparent Young’s modulus of the INL was significantly increased (Supplementary Figure 3c).

### 3.3 Ex vivo mechanical tissue properties are changing post mortem

In certain instances, the conduction of mechanical measurements on cells and tissues encounters limitations when attempted within a living organism. Factors such as limited optical penetration depth and the organ’s internal positioning may force the researcher to procure dissected or even sectioned tissues. *Ex vivo* samples, however, exhibit a reliance on a variety of artificial environmental conditions, such as media composition, pH or temperature. Notably, alterations in tissue architecture and composition that are intricately linked to the onset of decay processes may impact the read-out of mechanical measurements significantly. To evaluate the impact of these structural changes on mechanical properties measured through Brillouin microscopy and AFM-based indentation, we conducted timelapse measurements on identical specimens for a duration of up to eight hours *post mortem* (hpm) using the confocal Brillouin microscope (Figure 4*a*). Additionally, we conducted AFM-indentation measurements on retina sections that had been preserved to attain a comparable final time point. Our results show trends in mechanical properties of individual retinal layers for all quantified parameters. Brillouin microscopy revealed that the GCL and IPL remained constant in both the Brillouin frequency shift (Figure 4b) and linewidth (Figure 4c) within the first eight hours following the death of the organism. INL and IS/OS demonstrated subtle changes only at later time points. The OPL and ONL exhibited the most pronounced changes over time. Notably, all variations in the Brillouin frequency shift trended towards lower values, while the linewidth concomitantly appeared to change to increased values. None of these trends, however, displayed statistical significance. Complementary AFM-indentation experiments showed no statistically significant change of the apparent Young’s modulus after 1.5 hpm in comparison to 0.5 hpm. In the subsequent five hours *post mortem*, the resistance to compressive force had increased significantly only in the INL, but not in other layers (Figure 4d).

## 4. Discussion

AFM is currently considered the gold standard in quantitative mechanobiology^[3]^, due to its adaptability and comprehensible operation principle. It can operate within aqueous environments, allowing for precise control of a wide range of conditions *ex vivo*. Parameters, such as media composition, osmolarity, pH, temperature, humidity and CO_2_ concentration can be tightly regulated to probe living biological systems in appropriate and physiologically relevant settings^[40]^. Brillouin microscopy is currently still a nascent method in the field of biomedical and biomechanical research. Its optical, and therefore non-invasive, working principle, however, promises unprecedented mechanical insights into living organisms with an unparalleled combination of high spatial and temporal resolution. Because of its classification not only as an optical, but also as a mechanical method, and an obvious overlap of application with other mechanical testing methods, it has been consistently subjected to direct comparison with AFM-based measurements.

The presented study aimed at providing a systematic quantification of the mechanical properties of the murine retina using both Brillouin microscopy and AFM-based indentation with the intention to provide a basis not only for direct comparison, but also future studies on *ex vivo* retina tissues.

The mature retinal morphology is organized into three cell layers (the ganglion cell layer (GCL), the inner nuclear layer (INL), and the outer nuclear layer (ONL).These are separated by two synaptic layers, the inner plexiform (IPL) and the outer plexiform layer (OPL)^[41]^. The most distal retinal layer investigated in this work is situated adjacent to the ONL and contains the photoreceptor inner and out segments (IS/OS) ^[41]^. Our results show a mechanical profile of acute retina sections in which the Brillouin frequency shift displays an increase from the innermost layers, GCL and INL, towards the ONL, with a significant drop in absolute values in the IS/OS. The linewidth followed a similar, but less pronounced trend with significant differences only between the GCL and the ONL, the IPL and the ONL, as well as between the ONL and IS/OS. AFM-based indentation measurements yielded the comparatively highest values for the IPL, and a significant drop in absolute apparent Young’s moduli for the INL when compared to the neighboring IPL. Notably, the distinct Brillouin frequency shifts of the nuclear layers appear to reflect their difference with respect to cell body density and size, i.e. a small nuclear size with high density correlates to a high Brillouin frequency shift. Our findings are in line with previous studies on Brillouin microscopy measurement of retina tissues *ex vivo*^*[19]*^ and *in vivo*^*[42]*^. Our data furthermore corroborate reports showing that the nucleus has a higher Brillouin frequency shift as compared to other compartments of the cell^[19,22,43]^ leading to the assumption that the mere presence of a nucleus induces an increase in the Brillouin frequency shift, and that the number of nuclei would scale positively with an increase in the Brillouin frequency shift. The resistance to external loading as applied by AFM-based indentation, however, appears to be governed by different factors. Here, we detected no statistically significant difference among the individual nuclear layers. Since the estimated effect size is still large (see Methods section, *statistical analysis*), the lack of a statistical significance could be explained by an insufficient sample size. However, our current AFM-indentation results show comparable apparent Young’s moduli for all nuclear layers, but relative differences in their respective Brillouin frequency shift values. When compared to INL, for instance, GCL displays similar apparent Young’s moduli, but lower Brillouin frequency shifts. This mechanical discrepancy coincides with a difference not only in cell body density, but also a divergent expression of extracellular matrix (ECM) components^[44]^. The GCL has been shown to exhibit immunoreactivity to laminin, tenascin-C, and brevican which appear to be absent in the INL^[44]^. The specific compositional discrepancy between GCL and INL could induce a different mechanical phenotype when probed with either method. Additionally, this mechanical difference could depend on the time- and length-scale at which it is probed, and on additional mechanical and optical material properties that contribute to a distinct Brillouin frequency shift and an apparent Young’s modulus. However, the lack of a positive correlation between cell body density and the apparent Young’s modulus has been already demonstrated for zebrafish spinal cord tissue^[4]^ and could still be similarly true in murine retinae (this study). Our indentation measurements could identify a statistically significant difference between IPL and INL as well as between IPL and OPL. The plexiform layers are formed by reticulum of synaptic connections between different cell types and contain a multitude of extra cellular matrix (ECM) components^[44]^ that potentially dominate the mechanical tissue response on a time scale closer to that of indentation measurements. Interestingly, IPL and OPL differ significantly in their Brillouin frequency shifts and apparent Young’s moduli, but with an inverse respective relationship. This also points towards a divergent sensitivity of Brillouin microscopy and AFM-based indentation, possibly linked to the different length- and time-scales relevant for each method. The divergent behavior of the data sets obtained with Brillouin microscopy and AFM-based indentation is reflected in the lack of correlation between the two methods. While previous studies have found positive correlation between the Brillouin frequency shift and the apparent Young’s modulus^[21-23]^, our results show that there is no linear association between the variables in the different datasets. However, it does not necessarily imply the absence of a relationship between Brillouin frequency shift and apparent Young’s modulus. There could still be other types of more complex associations between the Brillouin frequency shift and the apparent Young’s modulus that are not captured by correlation coefficients. Moreover, Brillouin microscopy inherently involves the coupling of mechanical and optical material properties^[20]^. In fact, retinal cells exhibit extraordinary optical properties. Plexiform layers, for instance have been shown to cause light scattering, whereas Müller cells share optical, structural and geometric properties with optical fibers enabling low-scattering light guidance through the retinal layers^[45]^. Isolated retinal cells displayed comparable refractive indices in their somas which are lower than refractive indices of cellular structures residing in the IPL^[45]^. At the same time, the small nuclear size in murine retinae results in a higher average mass density of the central mass of highly refractive heterochromatin^[46]^. These optical characteristics could induce alterations in the mechanical profile of the retina tissue as measured with Brillouin microscopy, and could potentially establish a correlation between the longitudinal modulus and the apparent Young’s modulus. However, the direct determination of the refractive index distribution in extended, living tissues, along with the concurrent possibility of inferring the density from those measurements, is currently an unresolved challenge, despite considerable, recent advancements^[30]^. Likewise, the evaluation of AFM-indentation measurements to extract apparent Young’s moduli assumes either a constant Poisson’s ratio across all layers or combines it with the Young’s modulus as a fitting parameter that is extracted from force-distance curves. The exact determination of the Poisson’s ratio and its implementation in a direct comparison as presented in this study could contribute to the emergence of a correlation between the different moduli.

Our findings are supported by results we obtained by investigating the implemented measurement regiment to tissues that had been altered with genipin-mediated cross-linking and changes *post mortem*. After incubation with genipin, Brillouin frequency shifts approach significantly lower values in the GCL and the INL, but not in other layers. The same treatment caused an increase of the apparent Young’s modulus of the IPL while all other layers did not show a significant change upon genipin exposure. Genipin has been used to cross-link tissues and hydrogels while maintaining cell viability. In our hands, it evoked changes in two nuclear layers that were detectable by Brillouin microscopy, while AFM-indentation was sensitive to a change within the IPL. This further underscores the divergence of Brillouin microscopy and AFM-based indentation measurements. Interestingly, we found that time interval *post mortem* had similarly opposing effects. Brillouin frequency shift and linewidth displayed noticeable, but statistically nonsignificant mechanical changes over time. AFM-indentation showed a significant increase in values at six hours *post mortem* in the INL. Although not statistically significant, the observable trends for the Brillouin frequency shift and the apparent Young’s modulus follow opposing directions. However, the test statistic of the Kruskal-Wallis ANOVA employed for the data set obtained with Brillouin microscopy still contributed to a large effect size, which indicates a practical significance. This means that a true effect is likely concealed by the small sample size, and that our current sample size of three tissue samples per time point and layer does not allow to rule out the presence of decay-accompanying effects. Most importantly, these results point towards a decay-sensitive mechanical read-out that needs to be taken into account when designing future mechanical experiments on acute tissue sections. It also highlights the need for non-invasive, fast testing methods that could be employed *in vivo*.

## 5. Conclusion

Our integrative approach provides a comprehensive investigation of the distinct mechanical characteristics of individual retinal layers, directly comparing Brillouin microscopy with AFM-based indentation measurements. Given the considerable differences in sensitivities with respect to length- and time-scale, we argue that instead of employing both methods with the aim of one verifying the respective other, more in depth knowledge is to be gained when treating both methods as complementary. In fact, our results highlight that the *a priori* assumption of a correlation of the Brillouin frequency shift as measured with Brillouin microscopy and the apparent Young’s modulus as measured with AFM-based indentation measurements should be subjected to scientific scrutiny for each sample and experimental condition.

Future investigations might conduct an in-depth analysis of compositional and histological factors to connect a specific tissue architecture to a distinct mechanical output, focusing on both individual and synergistic effects. This approach would not only elucidate the nuanced alterations in mechanical properties observed in individual retinal layers, as reported in this study, but also lay the groundwork for identifying overarching principles applicable to diverse species, length scales, and time scales. By design, our results cannot serve to predict a mechanical profile of the murine retina that would occur inside a living organism. Leveraging optically transparent organisms in future analysis, however, could extent insights into the determinants of tissue mechanical properties *in vivo*. Such endeavors may facilitate the identification of common factors bridging the gap between experimental settings and hold promise for discerning universal principles governing tissue mechanics.

## Acknowledgements

We would like to thank all members of the Guck Lab for continuous support and valuable feedback throughout the development of the study. In particular, the authors gratefully acknowledge expert technical assistance from Raimund Schlüßler, Shada Abuhattum, and Eoghan O’Connell, and thank Jona Kayser for generously sharing microscopy equipment. We thank Timon Beck for critical review of the manuscript. The authors acknowledge financial support from the Deutsche Forschungsgemeinschaft (project number 460333672 – CRC1540 Exploring Brain Mechanics (subproject B03) to S.M. and J.G..

## Ethical statement

All experiments were planned and conducted under the principle of 4R’s: reduction, refinement, replacement and responsibility. Only surplus animals from regular breeding were used. The number of animals per experiment was reduced to the absolute minimum by using multiple samples from each animal and utilizing multiple measurement setups simultaneously. The performance of each technical setup was constantly monitored and optimized. All animals were kept under optimal conditions. The permit for animal sacrifice for the purpose of organ removal is filed under TS 13/2023 MPI at the animal welfare office of Friedrich-Alexander-Universität Erlangen-Nürnberg.

## Authors’ contributions

S.M., M.G., J.G., L.M.: conceptualization, project administration. S.M., M.G.: data acquisition, analysis, investigation, methodology, visualization. J.B.S.: data acquisition, analysis. S.M.: writing—original draft and writing—review and editing. P.M.: software. K.K., S.A.: methodology, microscopical quality assurance. J.G., L.M.: supervision, funding acquisition. All authors: review, editing.

All authors gave final approval for publication and agreed to be held accountable for the work performed therein.

---

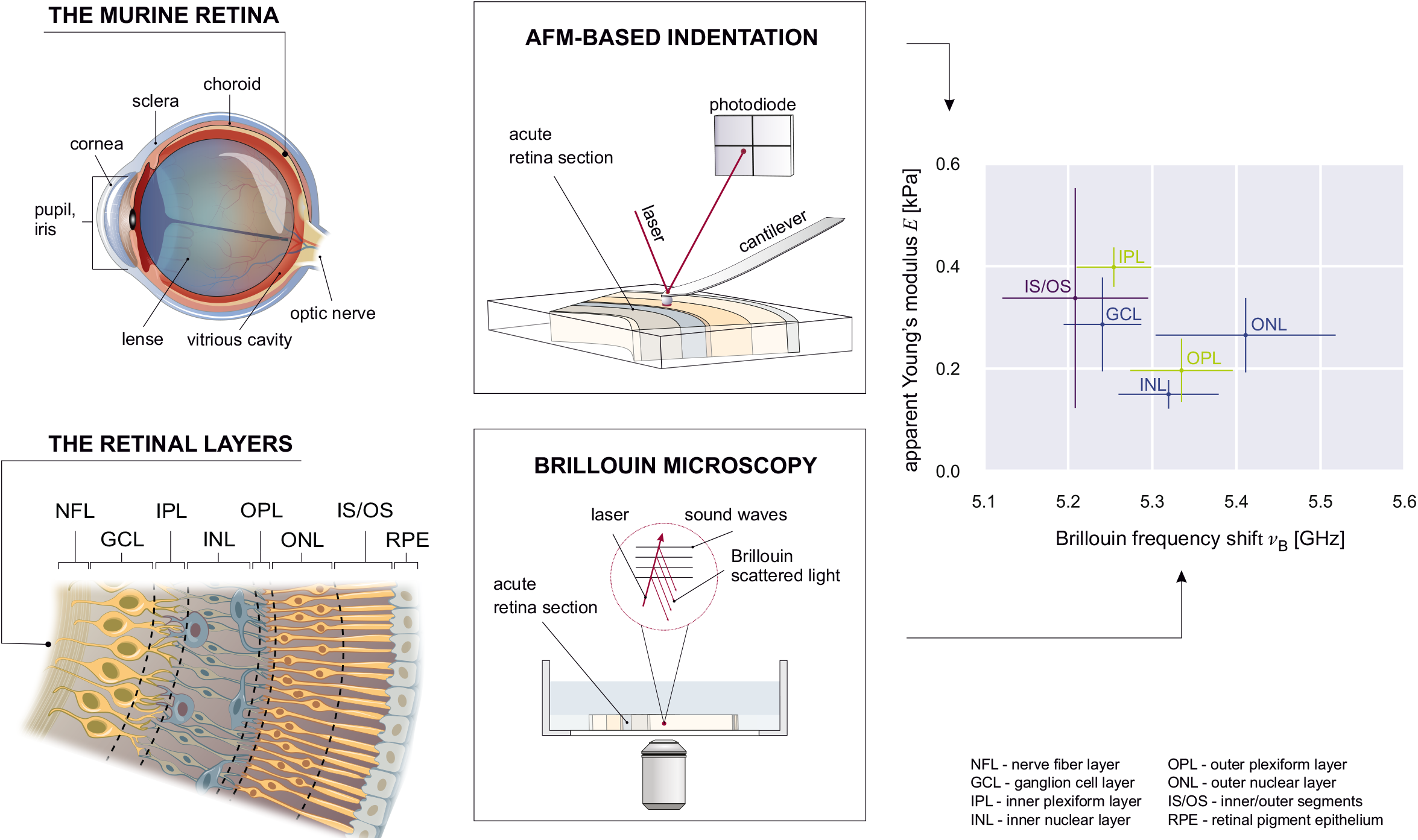

