## Supplemental Information for "Beyond comparison: Brillouin microscopy and AFM-based indentation reveal divergent insights into the mechanical profile of the murine retina"

Received xxxxxx
Accepted for publication xxxxxx
Published xxxxxx

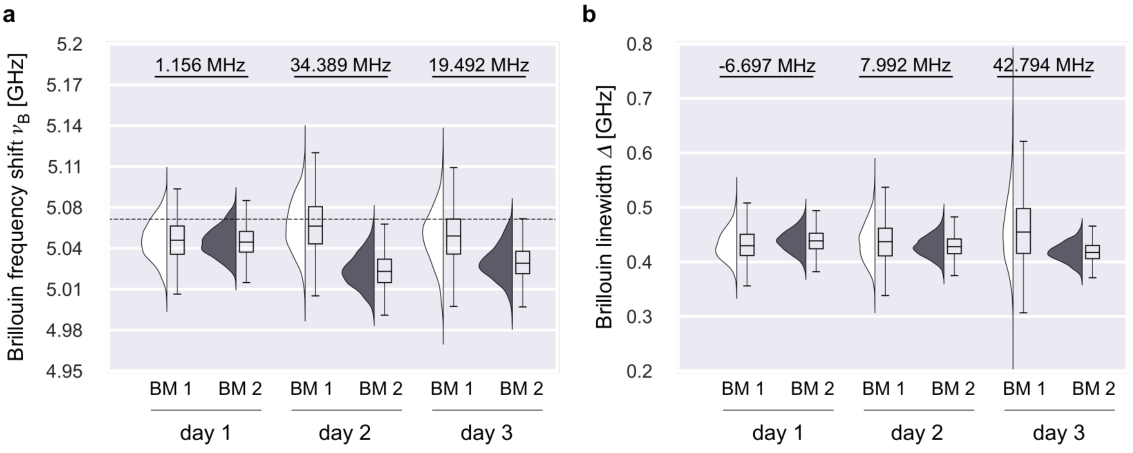


***Supplementary Figure 1: Water measurements for offset correction.*** **a)** Brillouin frequency shift with theoretical value for water at 5.066 GHz (dashed line), and **b)** linewidth for water obtained with the two setups BM 1 and BM 2 on three consecutive measurement days. Absolute differences of respective values are denoted above, and were added to values obtained with setup BM2. Graphs display raincloud plots with half violins showing the data distribution. Box plots showing the median, interquartile range (IQR) and whiskers for minimum and maximum data points. The scattered data points show all means including outliers (>1.5 IQR).

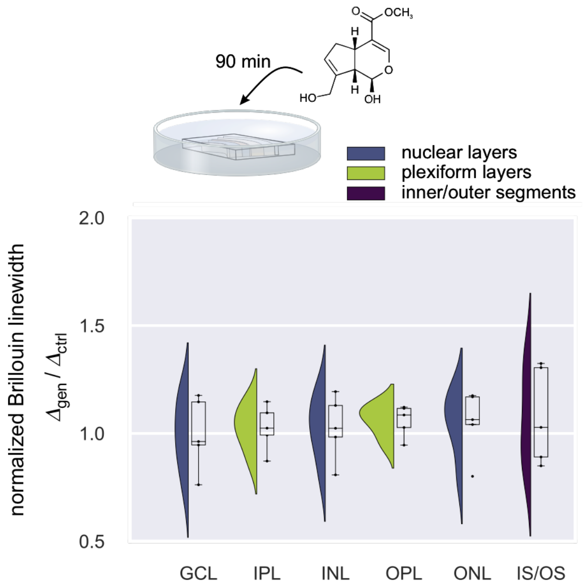


***Supplementary Figure 2: The impact of genipin cross-linking on the Brillouin linewidth of the retina***. The Brillouin linewidth of genipin treated sections (N=5) normalized to layer means from respective control measurements (N=5). Graphs display raincloud plots with half violins showing the data distribution. Box plots showing the median, interquartile range (IQR) and whiskers for minimum and maximum data points. The scattered data points show all means including outliers (>1.5 IQR). P-values indicate significance level of difference between retinal layers.

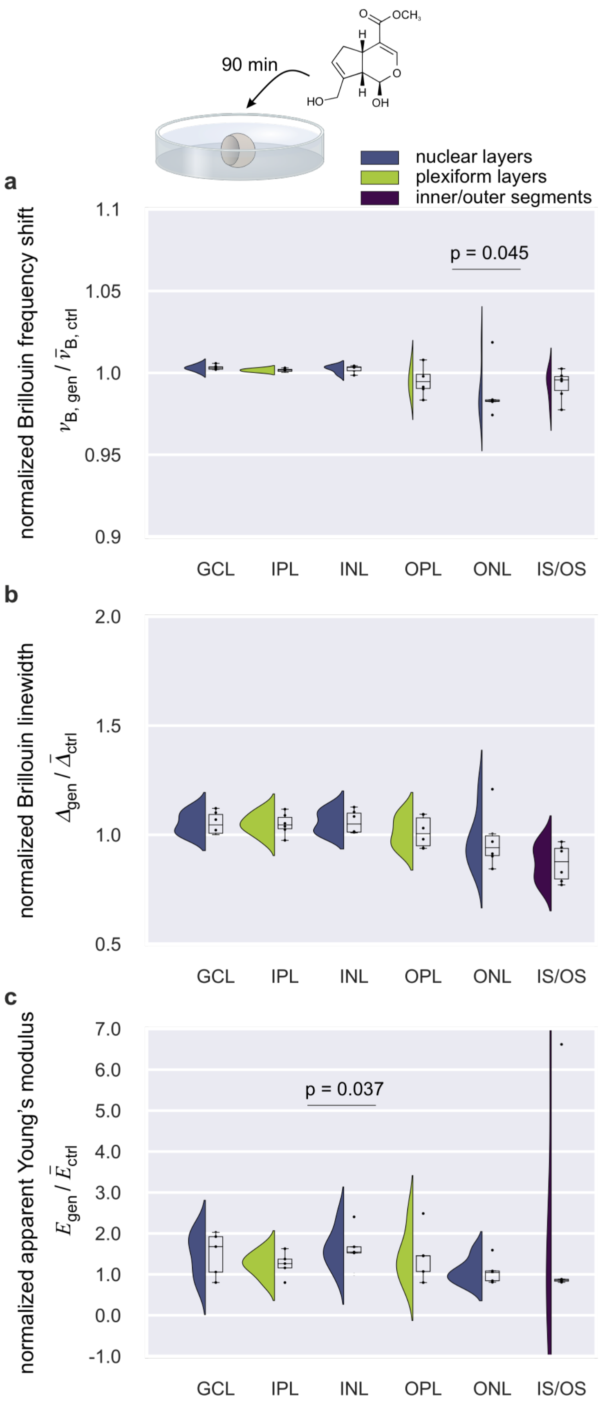


***Supplementary Figure 3: The impact of genipin cross-linking on the mechanical properties of the retina***. **a)** The Brillouin frequency shift (N=6), **b)** the Brillouin linewidth (N=6) and **c)** the apparent Young's modulus (N=5) of retinal sections after the retinal cup had been exposed to genipin. Values obtained from genipin treated cups were normalized to layer means from respective control measurements (N=6 for Brillouin microscopy and N=5 for AFM indentation). Graphs display raincloud plots with half violins showing the data distribution. Box plots showing the median, interquartile range (IQR) and whiskers for minimum and maximum data points. The scattered data points show all means including outliers (>1.5 IQR). P-values indicate significance level of difference between retinal layers. The pairwise comparisons using the Mann-Whitney test yielded for the ONL in a) U=31, Z=2.0016 which, with N=6, amounts to r=0.82 (large effect). For the INL in c) U=2, Z=-2.08893 which, with N=5, amounts to r=0.93 (large effect).
